# To praise or to blame? Neural signals predict divergent responses to moral hypocrisy

**DOI:** 10.1101/2022.07.25.501489

**Authors:** Jinting Liu, Jiamiao Yang, Fang Cui

**Author notes:** Nanhai Ave 3688, Shenzhen 518060, Guangdong, China. Contact. **Author Contributions** The first two authors contribute equally. J.L. and F.C. designed the study; J.Y. conducted the experiments and collected data; F.C., J.L., and J.Y. analyzed data; F.C., J.L., and J.Y. wrote the paper. **Declaration of ethics** All procedures performed in this study were in accordance with the 1964 Helsinki declaration and its later amendments or comparable ethical standards. The local ethics committee approved the experimental protocol. **Data and Code Availability Statement** The data and code are available through contacting the corresponding author at.

## Abstract

Moral hypocrisy is common in society and could be discouraged if observers always respond negatively. Understanding the observers’ responses to moral hypocrisy is critical for promoting genuine morality. This study took the perspective from the observers and compared their behavioral and neural responses to moral hypocrisy versus clear moral/immoral acts. Behaviorally, we found that claiming to help but avoiding the cost of help (i.e., moral hypocrisy) was endorsed as more moral than rejecting to help and even received monetary praise from 40% of observers. Neurally, moral hypocrisy versus clear moral acts elicited reduced responses in the reward system (e.g., ventromedial prefrontal cortex) and increased responses in regions responsible for disgust (e.g., anterior insula). The neural signals together with the interplay to the mentalizing network (e.g., temporoparietal junction) predicted divergent responses, even five months later. Our findings provide behavioral and neural accounts of how hypocrisy is perceived and why it prevails.

**Significance Statement:**

- From the perspective of a third-party observer, this study showed that moral hypocrisy is indeed deceptive and induces observers’ divergent responses of praise or blame.
- Claiming to help but avoiding the cost of help (i.e., moral hypocrisy) was considered more moral than rejecting to help and even received monetary praise from 40% of observers.
- Using fMRI, this study revealed that the divergent behavioral responses to moral hypocrisy could be predicted and classified by the activations of the reward system (e.g., ventromedial prefrontal cortex) and regions responsible for disgust (e.g., anterior insula) as well as the functional connectivity with the mentalizing network (e.g., right temporoparietal junction), even five months later.

## Introduction

Moral hypocrisy refers to the phenomenon that individuals appear moral while avoiding the costs of moral acts, such as claiming to help others but avoiding the cost of help, preaching one thing but practicing another, and judging others more harshly than judging themselves (Batson, Thompson, Seuferling, Whitney, & Strongman, 1999). Although hypocrites are despised by the general public, it is surprising that moral hypocrisy is very common in real life (Batson, Kobrynowicz, Dinnerstein, Kampf, & Wilson, 1997; Batson, Thompson, & Chen, 2002; Graham, Meindl, Loleva, Lyer R., & Johnson, 2015). As such, it has become an important topic in social and personality psychology (Batson et al., 2002; Graham et al., 2015).

Moral hypocrisy would be discouraged gradually if the observers always respond negatively. Therefore, understanding the observers’ responses to moral hypocrisy is critical for reducing hypocrisy and promoting genuine morality in society (Laurent, Clark, Walker, & Wiseman, 2014). However, most of the literature on moral hypocrisy takes the perspective of the actors, aiming to understand the motivations that drive the hypocritical behaviors (Batson et al., 1999; Chernyak, Harris, & Cordes, 2019; Dong, van Prooijen, & van Lange, 2019; Krettenauer, Bauer, & Sengsavang, 2019; Batson et al., 1997) and the factors that influence such behaviors (Batson et al., 1999; Caviola & Faulmuller, 2014; Carpenter & Marshall, 2009; Lammers, Stapel, & Galinsky, 2010). Little attention has been paid to the observers’ responses to moral hypocrisy.

By taking the perspective of the observers, this study aimed to understand how moral hypocrisy is detected, perceived, and judged by third-party observers at both behavioral and neural levels. Moral hypocrisy often occurs in the presence of behavioral inconsistency (Batson et al., 1997; Monin & Merritt, 2010). Thus, we designed a new paradigm to simulate situations with inconsistency between preceding claims and subsequent acts, in which the actors agreed to help a victim and then either transferred the cost of the help to others (i.e., intentional cost-free help which generates “moral hypocrisy”) or bore the cost themselves (i.e., intentional costly help which generates “clear moral act”).

For the observer, the actor’s preceding claim and subsequent act signal inconsistent information, thus introducing ambiguity in moral judgment (Batson et al., 1997; Bian, Li, Xia, & Fu, 2020). Some may blame the actor as a hypocrite and judge the act as immoral, while others may have different, even opposite, judgments (Caviola & Faulmuller, 2014). Once the inconsistency between claim and act is detected as moral hypocrisy, strong negative moral emotions, such as anger and disgust, would be triggered in the observers (Barden, Rucker, & Petty, 2005; Smith, Powell, Combs, & Schurtz, 2009). A behavioral study found that the stronger negative emotions experienced by the observers, the harsher punishments imposed on the hypocritical transgressors (Laurent et al., 2014). In real life, however, hypocrites usually succeed in deceiving the observers and gaining fame and fortune (Christen, 2014). If the observers fail to detect hypocrisy, they may praise the hypocrites as kind since their behaviors also signal moral motives by claiming their prosocial intentions. Stronger moral hypocrisy was found among individuals who had strong motives to gain good reputations (Dong, 2020). Taken together, considering the inconsistency between claims and acts, we hypothesized inconsistency of responses toward moral hypocrisy, i.e., greater variance and mixed valences in judging moral hypocrisy than clear moral acts.

Judgments of moral/immoral acts depend heavily on emotional reactions (Avramova & Inbar, 2013; Cummins & Cummins, 2012; Fourie, Thomas, Amodio, Warton, & Meintjes, 2014; Young & Koenigs, 2007). Observing immoral acts triggers aversive emotions such as anger and disgust (Vicario, 2016; Russell & Giner-Sorolla, 2013; Salerno & Peter-Hagene, 2013; Harenski & Hamann, 2006), while observing moral acts triggers rewarding emotions (Haidt J., 2003; Hu, Pornpattananangkul, & Nusslock, 2015). Brain regions involved in moral anger mainly include insula and amygdala (Harenski & Hamann, 2006). The anterior insula (AI), in particular, is also responsible for moral disgust (Boccadoro et al., 2021; Denke, Rotte, Heinze, & Schaefer, 2014; Ying et al., 2018; Zinchenko & Arsalidou, 2018). Immoral acts like unfair offers elicit the activation of AI (Sanfey, Rilling, Aronson, Nystrom, & Cohen, 2003), while moral acts like fair offers elicit the activations of the reward system such as ventromedial prefrontal cortex (vmPFC) and striatum (Tabibnia, Satpute, & Lieberman, 2008). Given these findings, we hypothesized that brain regions of negative emotions (e.g., AI and amygdala) would show greater responses to moral hypocrisy than to clear moral acts, while the reward system (e.g. vmPFC and striatum) would show the opposite effect.

Apart from emotional reactions, perception of the actor’s intention is critical for moral judgment as well (Cushman, 2008; Greene et al., 2009; Gan et al., 2015; Nobes, Panagiotaki, & Engelhardt, 2017). Recent fMRI studies on moral judgment have shown that people make spontaneous mental state inferences, as evidenced by the activation of brain regions responsible for theory of mind (Young, Cushman, Hauser, & Saxe, 2007; Young & Saxe, 2009). In moral hypocrisy, the inconsistency between claim and act does not provide clear clues about the actor’s intention, so moral judgment requires an inference of the intention (Isoda, 2016; Hesse et al., 2016). Therefore, we hypothesized that moral hypocrisy situations would activate the neural network of theory of mind (e.g., temporoparietal junction, posterior superior temporal sulcus, and medial prefrontal cortex) more strongly than clear moral acts (Bzdok et al., 2012; Schurz, Radua, Aichhorn, Richlan, & Perner, 2014). Further, the activations of brain regions of emotional reactions and theory of mind as well as their interactions would guide moral judgment so that the neural signals could predict the divergent responses toward moral hypocrisy.

## Materials and Methods

### Participants

The sample size commonly used in fMRI studies is 20-30 (Grady et al., 2021; Poldrack et al., 2017), which is considered insufficient for replication (Cremers et al., 2017; Grady et al., 2021; Turner et al., 2018). As such, this study aimed to recruit as many participants as possible within the constraints of our fundings. Forty-one right-handed participants were recruited from Shenzhen University. They were screened for a history of neurological disease, brain injury, and developmental disorders. All participants had normal or corrected-to-normal vision. Six participants were excluded from data analysis due to excessive head movements (rotation > 2°or translation > 2 mm; 5 participants) or misunderstanding of instructions (1 participant), leaving a final sample of 35 participants (19 females, mean age = 21.1 years, *SD* = 1.8). The study was conducted in accordance with the Declaration of Helsinki and was approved by the Ethical Committee of the Medical School, Shenzhen University. Written informed consent was obtained from each participant.

### Experimental Procedures

The fMRI experiment applied a 2 (Decider of cost: Helper vs. Computer) × 2 (Cost of help: Costfree vs. Costly) within-subject design. In the modified “third-party punishment paradigm”, participants were asked to observe the behaviors of the Helpers in a game and to morally judge them as good or bad. The game involved three parties: *Helper, Co-player*, and *Recipient*. At the beginning of each trial, a Helper decided whether to take two painful electrical shocks to help the Recipient exempt from 10 equally painful shocks. When the Helper decided to help, participants were then shown one of four conditions: Helper_Costfree, Helper_Costly, Computer_Costfree, or Computer_Costly. In the Helper_Costfree condition, the Helper decided to transfer the cost of help (i.e., 2 shocks) to the Co-player (i.e., moral hypocrisy) instead of bearing it her/himself, while in the Helper_Costly condition, the Helper decided to bear the cost of help by himself/herself (i.e., clear moral act). In the Computer_Costfree condition, the computer allocated the cost of help to the Co-player, while in the Computer_Costly condition, the computer allocated the cost of help to the Helper. After observing the behavior of the Helper, participants were asked to rate the morality of the behavior on a 9-point Likert scale (−4 = very immoral, 4 = very moral) and to decide whether to pay 1–4 MUs to reward or punish the Helper. They were given 4 MUs in each trial and each MU they paid would reduce or increase 2 MUs from the Helper’s final payoff. Figure 1A shows the timing and display of the task. There were 8 practice trials and 160 formal trials, with 32 trials for each condition (see Figure 1B for conditions), lasting about 1 h.

**Fig. 1.**
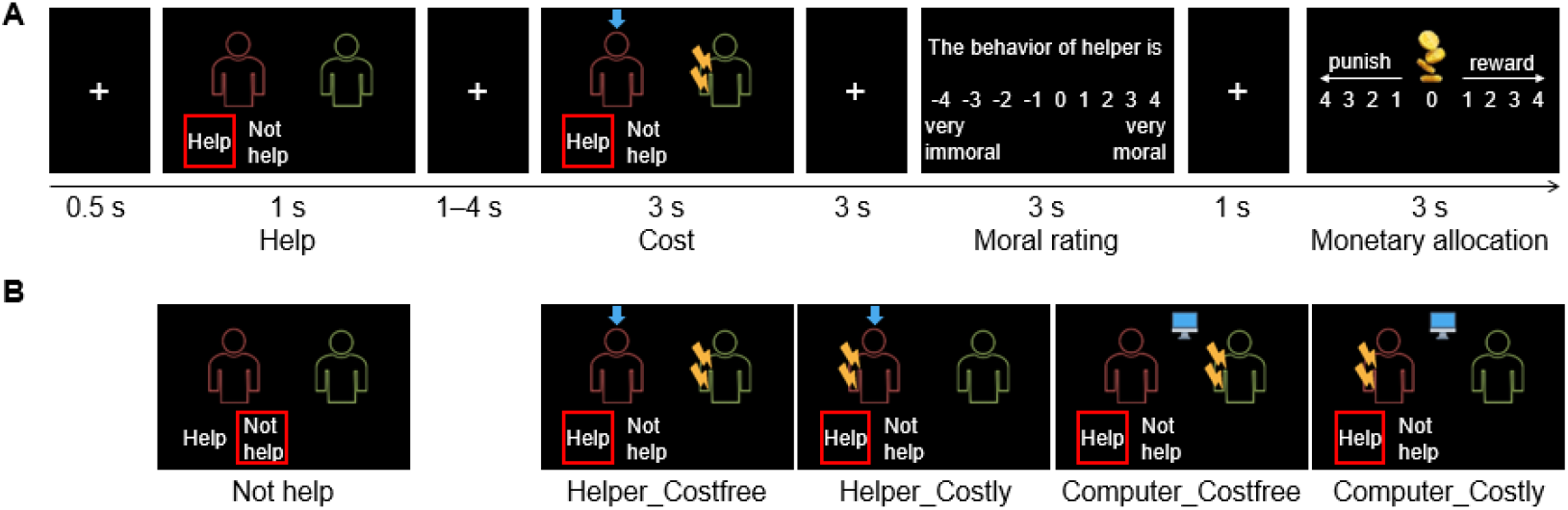
The timing, display, and design of the task. (A) Participants were told that there were three parties in the game: Helper (the red avatar), Co-player (the green avatar), and Recipient (not shown). On the Help Screen, participants saw whether the Helper decided to help the Recipient exempt from 10 painful shocks. The decision was highlighted by a red box. They then saw that either the Helper (the blue arrow icon pointing to the Helper) or the Computer (the computer icon) decided to allocate the cost of help (i.e., 2 shocks) to the Helper or the Co-player. In this example, the Helper decided to help the Recipient but transferred the cost of help to the Co-player (i.e., moral hypocrisy). Finally, the participants rated the morality of the Helper’s behavior and decided whether to pay 1–4 MUs to reward or punish the Helper. The starting points for the moral rating and monetary allocation were randomized. (B) The display of five conditions in the task. Of note, in the Not help condition, the Cost Screen and the following fixation were skipped.

The participants were instructed that (1) the behavioral data of the Helpers were randomly selected from a previous experiment; (2) in the previous experiment, each trial randomly selected three new strangers, randomly assigned the roles of Helper, Co-player, and Recipient to them, and randomly assigned the allocator of the cost of help to the Helper or the Computer; (3) the participants and the Helpers were paid according to the MUs in 10% of trials. Unbeknown to the participants, all players in the previous experiment were putative (see also Wang, Rao, & Zheng, 2017).

To examine whether neural signals could predict the judgments of moral hypocrisy in a long term, we invited the participants five months later and 33 participants responded to the questionnaire. They were shown each picture of Figure 1B and rated the morality of the Helpers’ behavior across five conditions four times on a 9-point Likert scale (−4 = very immoral, 4 = very moral).

### Image acquisition and preprocessing

We collected T2-weighted echo-planar images using a Siemens TrioTim 3.0T MRI machine at Shenzhen University. The images were acquired using a sequence with the following parameters: multiband = 4, repetition time = 1500 ms, echo time = 30 ms, flip angle = 75°, field of view = 192 × 192 mm^2^, number of slices = 72, slice thickness = 2 mm, voxel size = 2 × 2 × 2 mm^3^. We collected T1-weighted images using MPRAGE sequence (repetition time = 1900 ms, echo time = 2.23 ms, voxel size = 1.1 × 1.1 × 1.1 mm^3^).

Using SPM12 and DPABI, the images were slice-time corrected, motion-corrected, coregistered to the T1-weighted images (segmented with DARTEL algorithm; Nemoto, 2017), normalized to MNI space with a resolution of 2 × 2 × 2 mm^3^, smoothed with an isotropic 6-mm Gaussian kernel, and high-pass filtered at a cutoff of 128 s.

### Statistical Analysis

#### General linear modeling (GLM)

Statistical parametric maps were generated on a voxel-by-voxel basis with a hemodynamic model to estimate the neural responses of the Cost Screen (Figure 1A) at the individual level. The regressors of no interest entering into the general linear model (GLM) were onsets of every screen shown in Figure 1A. Six head-motion parameters were also included to control for the head motion nuisance. The regressors of interest were four conditions: Helper_Costfree, Helper_Costly, Computer_Costfree, and Computer_Costly. For individual-level analysis, the GLM with the four regressors were used to estimate the main effects and interaction effect of Decider and Cost. For group-level analysis, whole-brain one-sample *t*-tests were used to examine the effects. All results were corrected for multiple testing using thresholds of cluster-wise FDR-corrected *p* < .05 and voxel-wise uncorrected *p* < .001.

#### Region of interest (ROI) analysis

For illustration and further analysis, we extracted signals from regions of interest (ROIs) within a sphere of a 6-mm radius centered at the peak MNI coordinates. Three peak coordinates of whole-brain interaction (Table S2) were selected: vmPFC [−2 38 −18], left insula [−34 20 −4], and right insula [32 20 0].

To test the hypothesis that rewarding and aversive emotions triggered during the observation of clear moral acts and moral hypocrisy, we also extracted signals of a hypothesized ROI (vmPFC [−2 50 −6]) from a meta-analysis of neural correlates of the reward system (Bartra, McGuire, & Kable, 2013) and two hypothesized ROIs (left insula [−30 22 4] and right insula [38 22 −4]) from a meta-analysis of neural correlates of social norm violations (Boccadoro et al., 2021).

The extracted ROI signals were used to test the interaction effect of Decider and Cost on neural responses as well as the correlation between neural responses and moral ratings during scanning and five months later. To evaluate how well the neural signals correctly classified the praising and the blaming groups of moral hypocrisy, we created Receiver-Operating Characteristic (ROC) curves and used the area under the curve (AUC) as the accuracy of the classification. To evaluate the joint classification performance of multiple neural signals, we calculated a neural score using multiple logistic regression with multiple neural signals as predictor variables and the group as the outcome variable.

#### Psychophysiological Interaction (PPI) analysis

Given the role of the brain regions of emotional reactions in judging moral hypocrisy, a further question is: which regions would change their connectivity with the brain regions of emotional reactions during the observation of moral hypocrisy and clear moral acts? To address the question, we performed psychophysiological interactions (PPIs) analysis (Friston et al., 1997) using the vmPFC [−2 38 −18] and bilateral insula ([−34 20 −4]; [32 20 0]) identified in whole-brain interaction as seed regions. For each participant, the mean time series of each seed ROI (a sphere of 6-mm radius) were extracted and adjusted using *F*-contrast. The signals were high-pass filtered at a cutoff of 192 s to remove low-frequency drifts. Each GLM included the following PPI regressors: the psychological parameter (i.e., the contrast of (Helper_Costly − Helper_Costfree) > (Computer_Costly − Computer_Costfree)), the physiological activity (i.e., the interaction effect of ROI activity), and the interaction between the psychological parameter and physiological activity. Six head-motion parameters were also included as covariates to regress out any head-motion artifacts. The thresholds for multiple testing corrections were the same as previously reported.

## Results

### Behavioral results

We performed 2 (Decider: Helper vs. Computer) × 2 (Cost: Costfree vs. Costly) repeated ANOVAs on moral ratings and monetary allocations. We found significant main effects of Decider and Cost as well as their interaction, all *p*s < .008 on both moral ratings and monetary allocations, except the main effect of Decider on monetary allocations, *F*(1,34) = 2.29, *p* = .140. As shown in Figure 2A, participants rated costly help more morally than cost-free help (main effect of Cost: *F*(1,34) = 136.07, *p* < .001, partial η^*2*^ = .80), and this effect was more pronounced when the decision of cost was made by the Helper than the Computer (interaction: *F*(1,34) = 121.90, *p* < .001, partial η^*2*^ = .78). Likewise, for monetary allocations (Figure 2B), participants also rewarded the costly help more money than cost-free help (main effect of Cost: *F*(1,34) = 93.99, *p* < .001, partial η^*2*^ = .73), and this effect was more pronounced when the decision of cost was made by the Helper than the Computer (interaction: *F*(1,34) = 84.77, *p* < .001, partial η^*2*^ = .71).

**Fig. 2.**
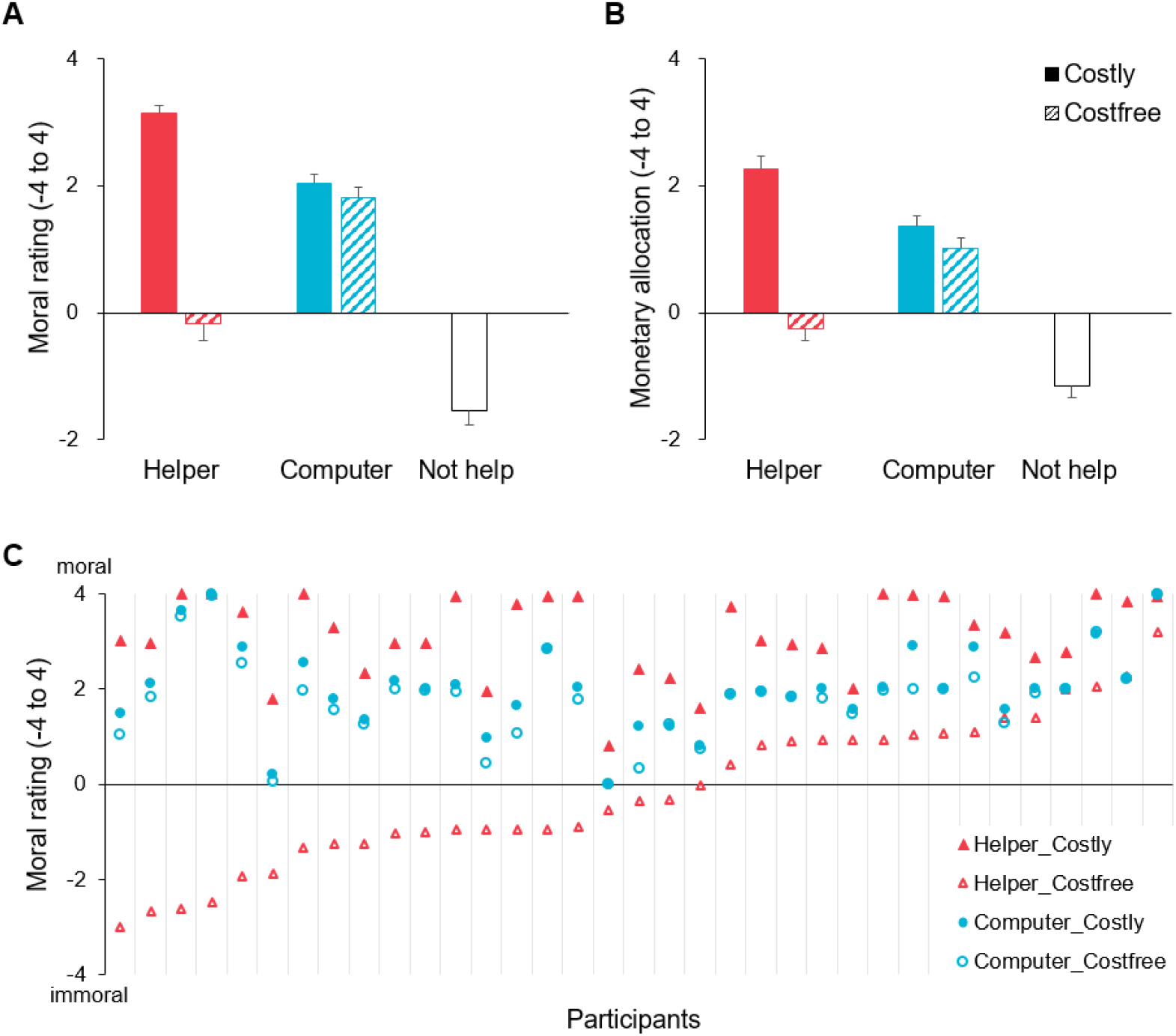
Behavioral results. (A) Mean moral rating as a function of Decider (Helper vs. Computer) and Cost (Costly vs. Costfree). Each pairwise comparison among the five bars was significant (*p*s < .001). (B) Mean monetary allocation as a function of Decider and Cost. Each pairwise comparison among the five bars was significant (*p*s < .002). (C) Mean moral rating as a function of Cost and Decider in each participant. For each participant, moral ratings were highest in Helper_Costly condition and lowest in Helper_Costfree condition. Twenty out of 35 participants rated the Helpers in Helper_Costfree condition as immoral (blaming group), while others judged the Helpers as moral (praising group).

As shown in Figures 2A and 2B, moral ratings and monetary allocations were highest in Helper_Costly condition, followed by Computer_Costly, Computer_Costfree, Helper_Costfree, and Not help conditions. Post-hoc paired *t*-tests revealed that each pairwise comparison of moral ratings among the five conditions was significant, all uncorrected *p*s < .001, so were those of monetary allocations, all uncorrected *p*s < .002. Considering the high within-subject correlations between moral ratings and monetary allocations (mean of *r*s = 0.83 ± 0.17) and the confounding of willingness to pay in monetary allocations, the following analyses focused on moral ratings.

As shown in Figure 2C, the rankings of moral ratings among conditions were fairly consistent across participants. However, the moral ratings of moral hypocrisy (i.e., Helper_Costfree condition) varied substantially across participants. *F* tests indicated a significant larger variance in Helper_Costfree condition (*s*^2^ = 2.42) than those in Helper_Costly (0.71; *F*(34, 34) = 3.43, *p* < .001), Computer_Costfree (0.89; *F*(34, 34) = 2.72, *p* = .004), and Computer_Costly (0.81; *F*(34, 34) = 2.98, *p* = .002) conditions. Moreover, moral ratings and monetary allocations in Helper_Costfree condition were not statistically different from zero, *p*s > .202, while those in Helper_Costly, Computer_Costfree, and Computer_Costly conditions were significantly larger than zero, all *p*s < .001.

### fMRI results

#### GLM

Contrasting brain activation during the observation of cost-free help versus costly help, we found significantly stronger activations in the left insula [−30 22 0] and the right insula [30 26 0] for Costfree > Costly, and stronger activation in the vmPFC [2 42 −18] for Costly > Costfree. Regarding the main effect of Decider, we found significantly stronger activation in the left insula [−30 20 −16] for Helper > Computer, and Lingual gyrus [2 −86 −10] for Computer > Helper. Regarding the interaction contrast of (Helper_Costly − Helper_Costfree) > (Computer_Costly − Computer_Costfree), we found significantly stronger activation in the vmPFC [−2 38 −18]; for the reversed interaction contrast, we found stronger activations in the left insula [−34 20 −4] and the right insula [32 20 0] (see Figure 3, Table S1, and Table S2 for comprehensive activations).

**Fig. 3.**
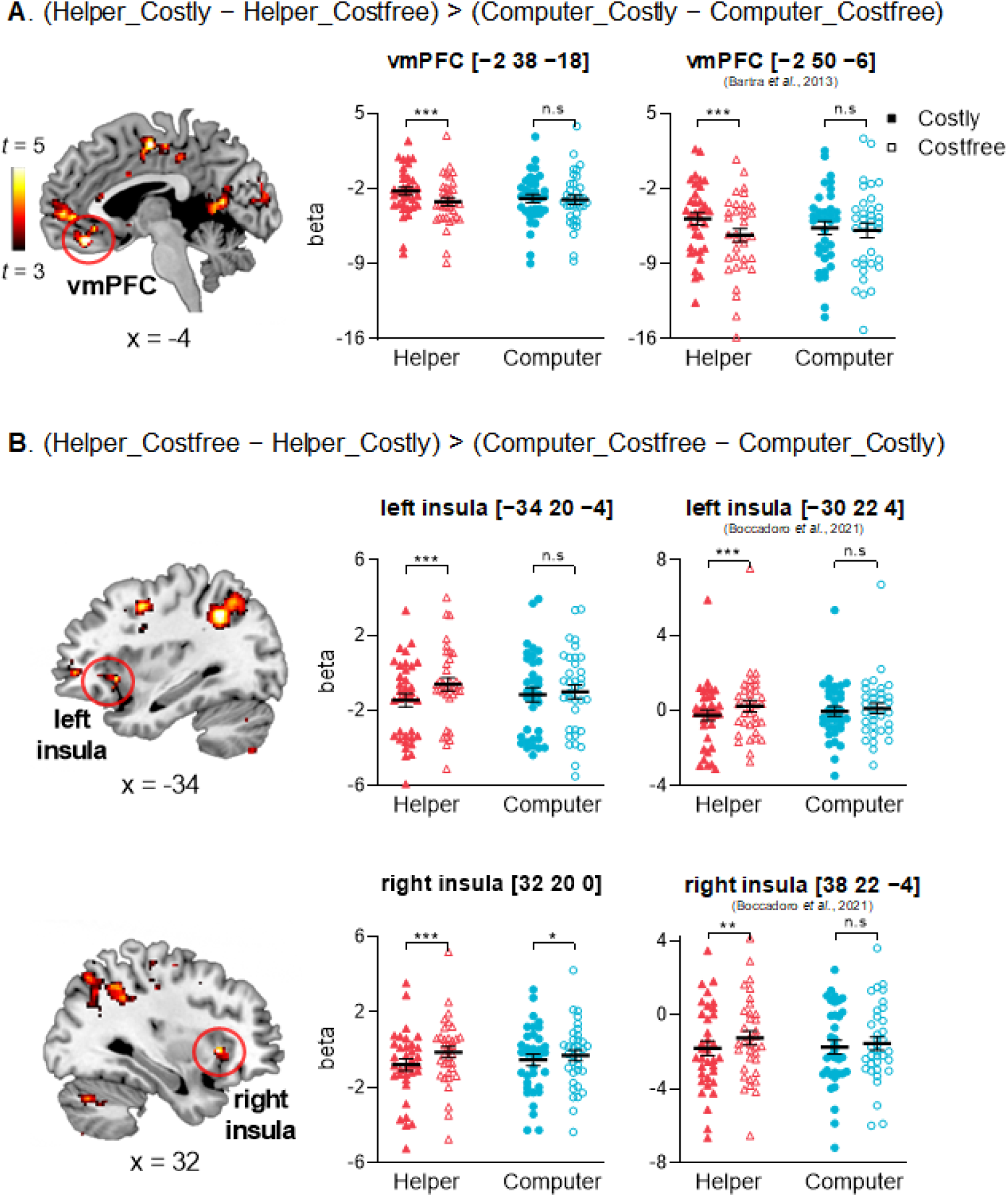
Whole-brain contrast and parameter estimates for the interaction effects. (A) A group-level parametric map of the contrast (Helper_Costly − Helper_Costfree) > (Computer_Costly − Computer_Costfree). Stronger activation in the vmPFC [−2 38 −18] was found during viewing costly help versus cost-free help and this main effect was more pronounced when the decision of cost was made by the Helper than the Computer. The main effect of Cost and the interaction were also found in the hypothesized ROI [−2 50 −6]. (B) A group-level parametric map of the contrast (Helper_Costfree − Helper_Costly) > (Computer_Costfree − Computer_Costly). Stronger activations in the left insula [−34 20 −4] and the right insula [32 20 0] were found during viewing cost-free help versus costly help and the main effects were more pronounced when the decision of cost was made by the Helper than the Computer. The main effects of Cost and the interactions were also found in the hypothesized ROIs of [−30 22 4] and [38 22 −4]. ROI signals extracted within a sphere of a 6-mm radius centered at the peak MNI coordinates.

#### ROI

The results of the three ROIs derived from our study confirmed the statistical inferences from the whole-brain GLM analysis. Figure 3A showed stronger activation of the vmPFC [−2 38 −18] during viewing costly help versus cost-free help, *F*(1, 34) = 29.55, *p <* .001, partial η^*2*^ = 0.47, and this main effect was more pronounced when the decision of cost was made by the Helper than the Computer, *F*(1, 34) = 30.36, *p <* .001, partial η^*2*^ = 0.47. On the contrary, Figure 3B showed stronger activations of the left insula [−34 20 −4] and the right insula [32 20 0] during viewing cost-free help versus costly help (*F*(1, 34) = 18.75, *p <* .001, partial η^*2*^ = 0.36; *F*(1, 34) = 26.47, *p <* .001, partial η^*2*^ = 0.44, respectively), and the main effects were more pronounced when the decision of cost was made by the Helper than the Computer (*F*(1, 34) = 19.86, *p <* .001, partial η^*2*^ = 0.37; *F*(1, 34) = 11.40, *p* = .002, partial η^*2*^ = 0.25, respectively).

The results of the three hypothesized ROIs showed similar patterns (Figure 3). The main effects of Cost and the interactions on activations of the vmPFC [−2 50 −6], the left insula [−30 22 4], and the right insula [38 22 −4] again reached significance, all *p*s < .045.

#### PPI

We found that the functional connectivity between the left insula [−34 20 −4] and the right temporoparietal junction changed significantly with the interaction contrast of (Helper_Costly − Helper_Costfree) > (Computer_Costly − Computer_Costfree), rTPJ [50 −48 10], *k* = 190, *t*(34) =5.21 (Figure 4C and Table S3). To verify the location of rTPJ, we conducted a PPI analysis within a functional mask of rTPJ (Bzdok et al., 2012), and the peak coordinate is almost unchanged, rTPJ [50 −46 10], *k* = 87, *t*(34) = 5.16. No significant result was found for the PPI analyses of vmPFC or right insula.

**Fig. 4.**
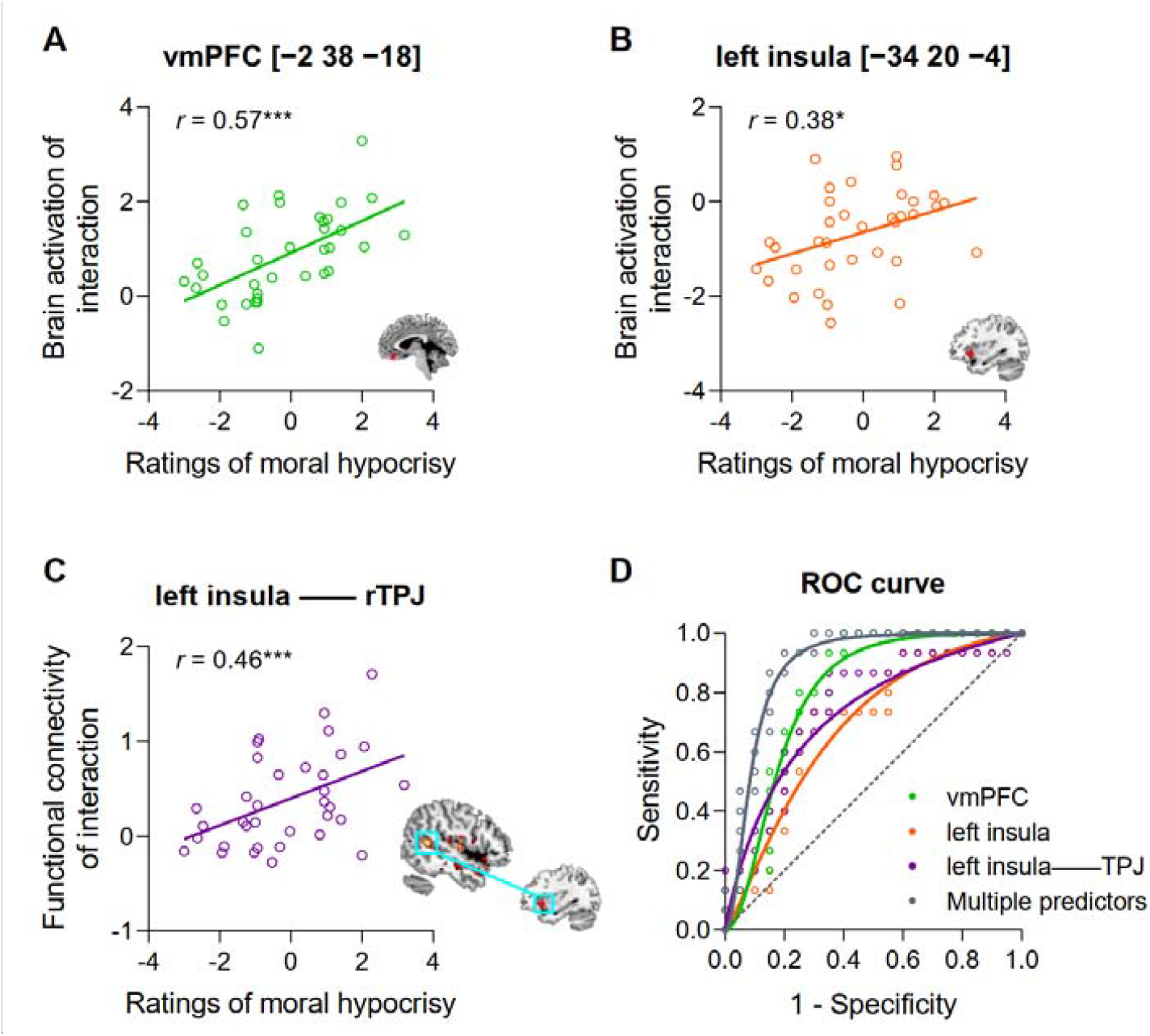
Predicting and classifying ratings of moral hypocrisy with the neural signals. (A) The interaction effect of vmPFC activation predicted the ratings of moral hypocrisy (i.e., Helper_Costfree condition). (B) The interaction effect of left insula activation predicted the ratings of moral hypocrisy. (C) The functional connectivity strength between the left insula and rTPJ predicted the ratings of moral hypocrisy. (D) The blaming group and praising group could be classified by the interaction effect of vmPFC activation, the interaction effect of left insula activation, and the functional connectivity strength between the left insula and rTPJ, both independently and jointly. The ROC curve fitting was performed through linear regression using the Littenberg and Moses linear model (Moses, Shapiro, & Littenberg, 1993) and the fitted formula was produced using Curve Fitting for Receiver Operator Characteristics (ROCs) Program (http://www.obg.cuhk.edu.hk/ResearchSupport/StatTools/CurveFitROC_Pgm.php).

#### Predicting ratings of moral hypocrisy during scanning and five months later

Correlational analyses revealed that the interaction effects ((Helper_Costly − Helper_Costfree) − (Computer_Costly − Computer_Costfree)) of ROIs signals predicted the ratings of moral hypocrpisy (i.e., Helper_Costfree) during scanning (vmPFC [−2 38 −18], *r* = 0.57, *p* < .001; left insula [−34 20 −4], *r* = 0.38, *p* = .023; Figures 4A and 4B). Likewise, the functional connectivity strength between left insula and rTPJ predicted the ratings of moral hypocrpisy during scanning as well, *r* = 0.46, *p* = .006 (Figure 4C).

Moreover, the neural signals of the interaction effects (marginal) significantly predicted long-term judgments of moral hypocrisy five months later, vmPFC [−2 38 −18], *r* = 0.30, *p* = .090; left insula [−34 20 −4], *r* = 0.35, *p* = .047.

#### Classifying praising and blaming groups by neural signals

As previously described, the ratings of moral hypocrisy varied substantially across participants. Twenty out of 35 participants (57%; blaming group) judged moral hypocrisy as immoral and 17 participants monetarily punished the Helper, while 15 participants (praising group) judged moral hypocrisy as moral and 14 participants monetarily rewarded the Helper.

As shown by the ROC curves in Figure 4D, the blaming group and praising group could be classified by the interaction effect of vmPFC with 81.0% (±7.5%) accuracy (*p* = .002), the interaction effect of left insula activation with 70.0% (±9.1%) accuracy (*p* = .046), or the functional connectivity strength between left insula and rTPJ with 75.0% (±8.6%) accuracy (*p* = .012) independently, and by the three predictors jointly with 90.0% (±5.4%) accuracy (*p* < .001).

## Discussion

Combining a modified “third-party punishment” paradigm with fMRI, we investigated observers’ behavioral and neural responses to moral hypocrisy in contrast to clear moral/immoral acts. Results support our hypotheses. Behaviorally, we found that observers endorsed moral hypocrisy (i.e., Helper_Costfree) as less moral than clear moral acts (i.e., Helper_Costly), but less immoral than no help (i.e., Not help). Neurally, moral hypocrisy versus clear moral acts elicited reduced responses in vmPFC and increased responses in the bilateral anterior insula. Importantly, we observed the largest individual differences with mixed valences in judging moral hypocrisy. The divergent responses of praising and blaming could be predicted and classified by the activations of vmPFC and anterior insula as well as the functional connectivity between anterior insula and rTPJ, even five months later.

The increased moral ratings of moral hypocrisy versus no help suggest the effectiveness of moral hypocrisy in social contexts. The helpers avoided the cost of help in both moral hypocrisy and no help conditions, while the former claimed a good intention to help and the latter did not. The increased moral evaluations suggest that simply claiming kindness can win the favor of observers. It explains from the observers’ perspective why moral hypocrisy is so prevalent and why individuals with strong reputation management motives behave more hypocritically (Dong et al., 2019). Notably, the victims in moral hypocrisy condition refrained from the 10 painful shocks whereas those in no help condition did not. The different outcomes for victims may contribute to the increased moral evaluations, as moral judgment is largely outcome-based (Young et al., 2010). Given the previous report of harsher punishments for hypocritical criminals than for non-hypocritical criminals (Laurent et al., 2014), future research needs to examine whether moral hypocrisy leaving victims to suffer would be judged as more immoral than no help.

The behavioral and neural changes between moral hypocrisy and clear moral acts cannot be attributed to cost alone, because these changes were much larger than those between accidental cost-free help and accidental costly help, as evidenced by the significant interactions. Comparing moral hypocrisy and clear moral acts, the former revealed an intention to avoid the cost of help while the latter did not. As a result, moral hypocrisy was judged as less moral than clear moral acts, accompanied by increased activations in anterior insula and reduced activations in vmPFC. Previous accounts of anterior insula in moral judgment include moral disgust towards norm violations (Denke et al., 2014; Harenski & Hamann, 2006; Ying et al., 2018; Moll et al., 2005) and empathic sadness for victims (Decety, Michalska, & Kinzler, 2012; Majdandzic, de, Feinberg, Aktar, & Bogels, 2012). Considering that moral hypocrisy is a violation of claimed moral standards (Batson et al., 1999) and frequently described as disgusting (Haidt, Rozin, McCauley, & Imada, 1997) and that the victims did not suffer from the shocks in both conditions, the increased responses in anterior insula may encode the feelings of disgust towards hypocritical agent rather than empathic responses for victims. Hypocrisy not only elicited negative feelings but also degraded the subjective values of the help, as reflected by the reduced responses in vmPFC. vmPFC is considered to integrate positive and negative reward signals into an abstract representation of subjective value of moral stimuli (Shenhav & Greene, 2010; 2014; Tabibnia, Lieberman, & Craske, 2008). The increased responses in vmPFC possibly encode high subjective value assigned to the authenticity in clear moral acts, as in the case of fair offers or saving lives (Shenhav & Greene, 2010; Tabibnia et al., 2008). The activation of vmPFC offers a neural account of why observing clear moral acts triggers rewarding emotions (Haidt J., 2003; Hu et al., 2015).

Parallel to a proposal that the inconsistency between claim and act brings ambiguity in moral judgment of hypocrisy (Batson et al., 1997; Bian et al., 2020; Monin & Merritt, 2010), we observed the largest individual differences with mixed valences in judging moral hypocrisy. Some praised it with moral rightness and monetary reward, while others blamed it with moral wrongness and monetary punishment. Activities in vmPFC and anterior insula tracked and classified the differential responses towards hypocrisy at that time and even five months later, suggesting that the divergent responses could be attributable to different processes of subjective value integration and aversive emotional reactions. In moral hypocrisy, unlike clear moral acts, the inconsistency between claim and act did not reveal intention clearly and required an intention attribution additionally (Isoda, 2016; Hesse et al., 2016). Some people may view the claim as a reputation-seeking behavior other than genuine kindness, whereas others may ignore the avoidance of help and consider the claim as moral. Intention attribution largely influences moral evaluation, by recruiting brain regions of theory of mind (Young et al., 2007; Young & Saxe, 2008; Young & Saxe, 2009). Indeed, functional connectivity between anterior insula and a typical region implicated in theory of mind (i.e., rTPJ) was associated with observers’ divergent responses toward moral hypocrisy, suggesting that rTPJ may guide observers to spontaneously represent the actors’ mental states underlying the inconsistency between claims and acts and then the interplay with anterior insula evokes aversive emotional reactions to perceived hypocritical intentions, which eventually makes observers blame the hypocritical acts.

Taken together, contrasting the observers’ behavioral and neural responses to hypocritical acts versus clear moral/immoral acts suggests that consistency in moral claims and acts is hedonically valued by recruiting brain regions of subjective value integration, whereas inconsistency underlying hypocritical acts recruits an interplay of brain regions implicated in spontaneous mentalizing and moral disgust, thereby leading observers to outrage. Nevertheless, simply claiming to help is considered less immoral than direct rejection and even receiving verbal and monetary praise from some observers. Our findings complement previous works from observers’ perspectives and provide behavioral and neural accounts of how hypocrisy is perceived and why it prevails.

## Acknowledgment

This work was supported by the National Natural Science Foundation of China (32171013 to F.C; 31600928 to J.L), Natural Science Foundation of Guangdong Province (2022A1515011097 to J.L), and the Science and Technology Innovation Commission of Shenzhen (JCYJ20210308103903001 to F.C).

---

**Table S1.** Results of whole-brain analysis of main effects

| Regions (Brodmann's area) | T | Cluster size (voxels) | MNI coordinates |  |  |
| --- | --- | --- | --- | --- | --- |
|  |  |  | X | Y | Z |
| Helper > Computer |  |  |  |  |  |
| L Superior frontal gyrus, medial part (9) | 6.10 | 175 | -6 | 58 | 34 |
| L Cerebellum | 5.91 | 131 | -18 | -80 | -38 |
| L Insula (47) | 4.70 | 75 | -30 | 20 | -16 |
| R Lingual gyrus (18) | 4.24 | 83 | 26 | -90 | -12 |
| L Inferior occipital gyrus (18) | 4.21 | 50 | -24 | -102 | -8 |
| Helper < Computer |  |  |  |  |  |
| R Lingual gyrus (17) | 5.77 | 325 | 2 | -86 | -10 |
| Costly > Costfree |  |  |  |  |  |
| L Rolandic operculum (48) | 7.94 | 54 | -44 | -2 | 10 |
| L Postcentral gyrus (3) | 7.40 | 899 | -36 | -22 | 48 |
| R vmPFC (11) | 6.76 | 1010 | 2 | 42 | -18 |
| L Inferior frontal gyrus, orbital part (47) | 5.52 | 93 | -32 | 40 | -16 |
| R Fusiform (19) | 5.30 | 322 | 22 | -62 | -12 |
| R Cuneus (18) | 4.79 | 48 | 14 | -90 | 30 |
| L Middle cingulate gyrus (23) | 4.45 | 34 | -6 | -10 | 46 |
| Costfree > Costly |  |  |  |  |  |
| L Precuneus (7) | 8.88 | 10297 | -6 | -74 | 38 |
| L Middle frontal gyrus (6) | 8.72 | 2380 | -28 | -4 | 52 |
| L Cerebellum | 8.31 | 4417 | -8 | -74 | -28 |
| R Cerebellum | 7.80 | 317 | 30 | -72 | -56 |
| L Supplementary motor area (32) | 7.57 | 1114 | 0 | 12 | 48 |
| R Insula (47) | 6.73 | 147 | 30 | 26 | 0 |
| L Cerebellum | 6.46 | 321 | -36 | -72 | -54 |
| L Cerebellum | 5.67 | 84 | -34 | -46 | -50 |
| L Insula (47) | 5.50 | 140 | -30 | 22 | 0 |
| L Cerebellum | 5.40 | 100 | -12 | -64 | -48 |
| R Calcarine (17) | 5.22 | 77 | 10 | -88 | 4 |
| R Vermis | 5.17 | 34 | 0 | -60 | -42 |
| L Inferior frontal gyrus, triangular part (45) | 4.86 | 122 | -54 | 16 | 0 |
| R Inferior temporal gyrus (20) | 4.74 | 81 | 54 | -48 | -16 |
| R Inferior frontal gyrus, orbital part (47) | 4.68 | 75 | 46 | 42 | -18 |
| L Superior temporal gyrus (22) | 4.68 | 108 | -64 | -44 | 16 |
| R Middle temporal gyrus (22) | 4.64 | 47 | 52 | -54 | 22 |
| L Middle frontal gyrus, orbital part (46) | 4.58 | 124 | -46 | 50 | -4 |
| R Heschl (48) | 4.55 | 59 | 42 | -20 | 12 |
| L Middle cingulate gyrus (23) | 4.34 | 33 | -6 | -26 | 24 |
| L Lingual gyrus (17) | 4.19 | 42 | -10 | -70 | 6 |

*Note:* Results are thresholded with voxel-wise uncorrected $p < .001$ and cluster-wise602 FDR-corrected $p < .05$ .

**Table S2.** Results of whole-brain analysis of interaction effect

| Regions (Brodmann's area) | T | Cluster size (voxels) | MNI coordinates |  |  |
| --- | --- | --- | --- | --- | --- |
|  |  |  | X | Y | Z |
| (Helper_Costly – Helper_Costfree) > (Computer_Costly – Computer_Costfree) |  |  |  |  |  |
| L Postcentral gyrus (4) | 9.42 | 1573 | –48 | –18 | 50 |
| R Cerebellum (37) | 9.20 | 1100 | 22 | –52 | –22 |
| L vmPFC (11) | 7.34 | 730 | –2 | 38 | –18 |
| L Calarine (17) | 5.43 | 169 | –10 | –56 | 8 |
| L Supplementary motor area (6) | 5.34 | 79 | –2 | –10 | 54 |
| R Superior occipital gyrus (19) | 5.10 | 88 | 16 | –88 | 32 |
| L Superior temporal gyrus (48) | 5.06 | 241 | –54 | –24 | 14 |
| L Anterior cingulate gyrus (24) | 4.82 | 98 | –2 | 28 | 14 |
| R Cerebellum | 4.37 | 44 | 16 | –62 | –48 |
| L Cuneus (18) | 4.34 | 50 | –6 | –90 | 26 |
| (Helper_Costfree – Helper_Costly) > (Computer_Costfree – Computer_Costly) |  |  |  |  |  |
| L Inferior frontal gyrus (48) | 5.90 | 859 | –52 | 18 | 16 |
| L Middle frontal gyrus, orbital part (47) | 5.89 | 341 | –40 | 50 | –4 |
| R Cerebellum | 5.85 | 438 | 26 | –66 | –32 |
| R Superior parietal gyrus (7) | 5.62 | 258 | 18 | –70 | 52 |
| L Inferior parietal gyrus (40) | 5.54 | 820 | –44 | –50 | 44 |
| L Supplementary motor area (8) | 5.19 | 407 | –4 | 18 | 52 |
| R Insula (47) | 5.14 | 37 | 32 | 20 | 0 |
| R Inferior frontal gyrus, triangular part (45) | 4.90 | 213 | 52 | 28 | 26 |
| R Angular (7) | 4.82 | 381 | 36 | –64 | 46 |
| R Middle frontal gyrus (9) | 4.75 | 90 | 44 | 12 | 52 |
| L Insula (47) | 4.63 | 47 | –34 | 20 | –4 |
| L Superior frontal gyrus (6) | 4.36 | 59 | –26 | –6 | 54 |
| L Lingual gyrus (18) | 4.29 | 54 | –14 | –86 | –14 |

*Note:* Results are thresholded with voxel-wise uncorrected $p < .001$ and cluster-wise FDR-corrected $p$ $< .05$ .

**Table S3.** Results of psychophysiological interaction analysis

| Regions (Brodmann's area) | T | Cluster size (voxels) | MNI coordinates |  |  |
| --- | --- | --- | --- | --- | --- |
|  |  |  | X | Y | Z |
| Seed 1: vmPFC [-2 38 -18] |  |  |  |  |  |
| none |  |  |  |  |  |
| Seed 2: right insula [32 20 0] |  |  |  |  |  |
| none |  |  |  |  |  |
| Seed 3: left insula [-34 20 -4] |  |  |  |  |  |
| R Precuneus (5) | 5.67 | 131 | 6 | -46 | 50 |
| R Temporoparietal junction (21) | 5.21 | 190 | 50 | -48 | 10 |
| L Paracentral lobule (4) | 5.15 | 71 | -12 | -34 | 66 |
| R Middle temporal gyrus (20) | 5.09 | 36 | 52 | -10 | -20 |
| L Calcarine (17) | 4.79 | 66 | -8 | -72 | 16 |
| L Superior temporal gyrus (48) | 4.76 | 39 | -56 | -14 | 6 |
| R Supplementary motor area (32) | 4.71 | 35 | 2 | 8 | 48 |
| R Supplementary motor area (6) | 4.63 | 50 | 12 | 4 | 56 |
| L Superior frontal gyrus (6) | 4.59 | 43 | -26 | -8 | 54 |
| R Rolandic operculum (48) | 4.58 | 47 | 56 | -18 | 20 |
| R Supplementary motor area (6) | 4.55 | 89 | 6 | -8 | 66 |
| L Superior temporal gyrus (42) | 4.40 | 45 | -62 | -42 | 16 |
| R Precentral gyrus (6) | 4.38 | 39 | 42 | -4 | 52 |
| L Superior occipital gyrus (19) | 4.35 | 33 | -24 | -78 | 34 |
| L Superior temporal gyrus (42) | 4.20 | 58 | -62 | -22 | 14 |
| L Inferior parietal gyrus (40) | 3.97 | 37 | -38 | -40 | 52 |

*Note:* Table S3 lists the PPI results of the interaction contrast of (Helper\_Costly – Helper\_Costfree) > (Computer\_Costly – Computer\_Costfree). Results are thresholded with voxel-wise uncorrected $p$ < .001 and cluster-wise FDR-corrected $p$ < .05. No cluster survived the threshold in the reverse interaction contrast, i.e., (Helper\_Costfree – Helper\_Costly) > (Computer\_Costfree – Computer\_Costly).

